# Cell type-specific associations with Alzheimer’s Disease conserved across racial and ethnic groups

**DOI:** 10.1101/2025.04.14.648597

**Authors:** Tain Luquez, Jonathan Algoo, Rebecca Chiu, Jason A. Mares, Archana Yadav, Matti Lam, Pallavi Gaur, Xiaoying Lai, Dylan I. Lee, Fahad Paryani, Rafe Batchelor, Irla Belli, Jordan Henry, Courteney Mattison, Lindsey Starr, Philip L. De Jager, Lisa L. Barnes, David X. Marquez, Ya Zhang, David A. Bennett, Vilas Menon

**Affiliations:** Center for Translational and Computational Neuroimmunology, Department of Neurology, Columbia University Irving Medical Center, New York, NY; Department of Kinesiology and Nutrition, University of Illinois at Chicago, Chicago, IL; Rush Alzheimer’s Disease Center, Rush University Medical Center, Chicago, IL

## Abstract

Genomic studies at single-cell resolution have implicated multiple cell types associated with clinical and pathological traits in Alzheimer’s Disease (AD), but have not examined common features across broad, multi-ethnic populations, and across multiple regions. To bridge this gap, we performed single-nucleus RNA-seq and ATAC-seq profiling of cortical and subcortical brain regions from post-mortem samples across Non-Latin White, African American, and Latin donors (the latter of any race). Using discrete and continuous dissection of molecular programs, we elucidate cell-type-specific glial and neuronal signatures associated with AD across multiple population groups. Notably, we found that multiple microglial (*GPNMB*+, *CD74*+, and *CR1*+ subgroups) and astrocyte (*SERPINH1*+ and *WIF1*+ subgroups) signatures are associated with worse clinical and pathological phenotypes across all three population groups. We also report continuous gene expression factors in oligodendrocytes that are not captured by discrete clusters, yet still show strong associations with disease phenotypes. Finally, we observe these discrete cellular identities and continuous gene programs separate cognitively impaired donors into 6 molecularly distinct subgroups that span racial and ethnic population groups. Overall, our study identifies key cell types and gene programs implicated in AD that are shared across population groups, and provides an initial data set that underscores how representative sampling can capture conserved signatures as well as disease heterogeneity, leading to better prioritization of key cell types for further investigation.

## Introduction

Alzheimer’s Disease (AD), the most common cause of dementia in older individuals, has a complex etiology resulting in neurodegeneration and memory loss. Studies of post-mortem human tissue have identified amyloid-beta aggregates and hyperphosphorylated tau tangles as the major pathologies associated with the disease, leading to a hypothesized cascade of pathology spreading, cellular alterations, circuit dysregulation, and cognitive impairment[1]. Although the ultimate cellular phenotype is degeneration and loss of neurons, transcriptomics and proteomics studies at the bulk[2–10] and single-cell level[11–24] have shed light on how almost all major cell classes in the brain appear to be altered in disease. However, most of these studies have profiled human tissue from individuals of European descent; given differences in prevalence of AD across racial and ethnic population groups[25], it is not immediately apparent whether previously reported cell type-specific disease associations are generalizable different population groups. In addition, only a handful of studies have explored multiple brain regions across large numbers of individuals to understand the extent of inter-regional difference in AD associations[20]. Finally, profiling an overall population of individuals with greater diversity in genetics and life exposure allows for better characterization of the molecular heterogeneity associated with the disease. In light of these gaps, we generated joint single-nucleus RNA and Assay for Transposase-Accessible Chromatin (ATAC)-sequencing data in three brain regions from 160 individuals who self-identified as African-American, Latin White, or non-Latin White, all of whom were participants in longitudinal studies with prospective brain autopsy based at the RUSH University Alzheimer’s Disease Center: the Religious Orders Study, Memory and Aging Cohort (ROSMAP)[26], Minority Aging Research Study (MARS)[27], and Latino Core[28] studies. We elected to address two questions with this data set: 1) which cell type-specific signatures are found consistently associated with disease in all three population groups, and 2) which molecular subtypes of disease may exist across this multi-ethnic set of individuals. Importantly, we focus on associations with self-reported race and ethnicity, rather than genetic ancestry, in order to include effects of life experience on brain signatures. As described below, we identified a subset of cell type-specific signals – mostly in microglia, astrocytes, and oligodendrocytes – associated with AD phenotypes in all three population groups, with differential impact in each brain region. Some, but not all, of these signatures have been implicated in prior studies[29]. In addition, our more representative population sampling identifes three transcriptomically-defined groups each among individuals with mild cognitive impairment and with dementia, building on prior evidence of molecular heterogeneity in aged individuals with cognitive decline. Overall, this study prioritizes a set of candidate cell types and gene signatures that are altered in broad and specific subpopulations in AD.

## Results

### Cell type overview and regional specificity in neurons and subsets of astrocytes

To identify differences in individuals with AD-related pathology and/or cognitive impairment relative to those without, we performed joint single-nucleus RNA- and ATAC-seq on nuclei from 399 samples, deriving from 56 self-identified Non-Latin African-American participants, 31 self-identified Latin participants (of any race), and 80 self-identified White participants (Fig. 1A,B). Three brain regions – dorsolateral prefrontal cortex (DLPFC), superior temporal gyrus (STG), and anterior caudate (AC) – were profiled from 94 individuals, with the remaining individuals having one or two regions profiled (Fig. 1C) due to tissue availability. Nuclear suspensions from three individuals were pooled prior to single-nucleus reverse transcription and library construction; each 3-sample pool was designed to include multiple brain regions and both sexes as far as possible, and nuclei were demultiplexed and assigned to donors using genetic data (See Data Availability statement for details on samples in each pool). After ambient RNA subtraction and quality control filtering, we iteratively clustered ∼1.9 million nuclei using RNA-seq data (Methods). Briefly, the first clustering round separated the major cell type classes, and three subsequent rounds of clustering identified groups of nuclei comprising doublets (based on mixed genetic and/or gene expression signatures). The final round of clustering identified expected and novel subpopulations within each major class, with the optimal number of subclusters determined by two clustering robustness metrics (Methods, Fig. 1D, S1, S2). The final set of nuclei had a median of 2,156 genes and 5,149 unique molecular identifiers (transcripts) detected, with some variation across major cell classes (Fig. S1). We did not find further subdivisions among these transcriptomic subclusters based on ATAC-seq data. This de novo clustering approach allowed us to generate a unified set of clusters across all three of our brain regions, some of which (e.g. AC) do not currently have published taxonomies suitable as references for label transfer.

**Figure 1:**
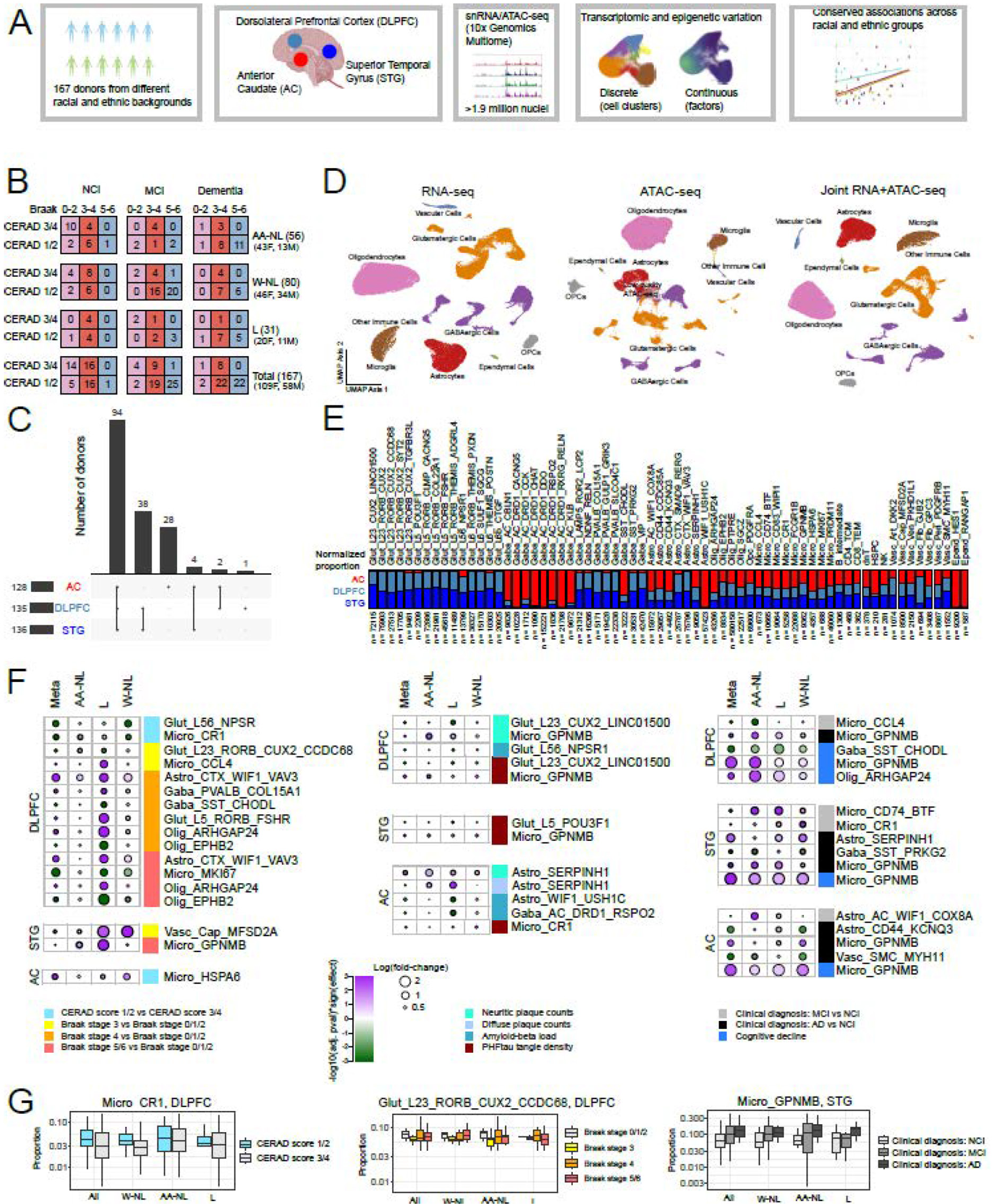
Discrete cell clusters and region-specific conserved associations with clinical and pathological traits across multiple population groups. A) Overview of experimental and analytical workflow of this study. B) Breakdown of pathological stages, cognitive diagnosis, and self-reported racial/ethnic groups of individuals in this study. AA-NL: Non-Latin African-American, L: Latin, W-NL: Non-Latin White. C) Upset plot showing number of individuals with combinations of brain regions profiled. AC: Anterior Caudate, DLPFC: Dorsolateral Prefrontal Cortex, STG: Superior Temporal Gyrus of the cortex. D) UMAP visualization of all (∼1.9 million) post-QC nuclei based on RNA-seq expression (left), ATAC-seq counts (middle), and joint RNA+ATAC-seq data (right). Major cell types are clearly identifiable visually in all three analyses. E) Relative abundance of each cell type subcluster across each of the three regions (double-normalized to sum to 1 for each cluster). Glutamatergic neurons are mostly confined to the cortical regions, whereas GABAergic neurons are grouped into distinct caudate and cortex-specific clusters. Some astrocyte clusters also show region-specificity, whereas oligodendrocytes and microglia are found across all regions. F) Summary of p-values and effect sizes for all subclusters with statistically significant (meta-analysis FDR-adjusted p-value <0.05 & I^2^<50%) and consistent associations with pathological and cognitive phenotypes across all three population groups, as shown in the legend. Effect sizes and FDR-adjusted p-values are from the ANCOM-BC method to assess cluster proportion associations. A subset of microglia and astrocyte subclusters show consistent associations across pathological and cognitive measures. NCI: Non-cognitively impaired, MCI: Clinical diagnosis of Mild cognitive Impairment, AD: Clinical diagnosis of Alzheimer’s dementia. G) Distributions of specific clusters showing associations with AD phenotypes. Left: The CR1+ microglia cluster is consistently lower in DLPFC from individuals with CERAD neuritic plaque scores 3 and 4 versus individuals with scores of 1 and 2. Middle: The RORB+/CUX2+/CCDC68+ glutamatergic neuronal cluster is lower in STG in individuals at Braak stage 3 and higher, as compared to individuals at Braak stage 0/1/2. Right: The GPNMB+ microglia cluster is more abundant in individuals with a clinical diagnosis of MCI or AD.

As expected, neuronal clusters showed region specificity, with glutamatergic neurons being primarily found in the two cortical regions (STG and DLPFC), and GABAergic neuronal clusters showing minimal overlap between the cortex and caudate, except for one neuronal subclass marked by somatostatin (*SST*) and chondrolectin (*CHODL*) expression, which was found in all three brain regions (Fig. 1E). Among glial clusters, only two classes showed any degree of regional specificity (Fig. 1E): ependymal cells (found primarily in the caudate) and astrocytes (with *SMAD9+/RERG+* and *WIF1+/VAV3+* astrocytes in the cortex and *WIF1+/USH1C+* astrocytes in the caudate). Overall, we identified 61 clusters across all major classes (Fig. S2), not including non-microglial immune cells, whose numbers were too low for robust clustering and were thus subject to label transfer using the Azimuth reference resource[30] (Methods).

Our primary goal was to identify cell subgroups that were consistently differentially abundant in relation to AD phenotypes across all three population groups. We examined discrete and continuous measures of AD phenotypes including CERAD neuritic plaque scores, Braak stages, neuritic plaque counts, diffuse plaque counts, amyloid-beta load, and PHFtau tangle density, as well as clinical diagnoses of No cognitive impairment (NCI), mild cognitive impairment (MCI), or Alzheimer’s dementia, and the slope of cognitive decline[26] (Methods), accounting for age at death, sex, and post-mortem interval as covariates, and education as an additional covariate for cognitive measures. To ensure robustness across all three population groups, we used a stratified approach whereby we assessed cluster proportion associations in each group and then performed a meta-analysis across the three groups, keeping only results that had an FDR-adjusted p-value <0.05, consistent sign of association in all three groups, and I^2^ < 50% (Methods, Supplemental Table 1). It is important to note that this stratified approach identifies conserved signatures associated with AD phenotypes despite our three population groups not being matched across demographic variables, most notably the age at death (Fig. S1).

### *GPNMB+*, *CD74*-high, and *CCL4+* microglial subgroups show consistent associations with AD traits across populations

The strongest consistent signal across population groups was the higher proportion of *GPNMB+* microglia in individuals with a clinical diagnosis of Alzheimer’s dementia and/or steeper slope of cognitive decline; this signal was found in all three brain regions (Fig. 1F). Higher proportions of this microglial subgroup were also associated with higher PHFtau tangle density in both cortical regions, and with higher neuritic plaque counts in the DLPFC; this latter finding is consistent with prior single-nucleus studies on DLPFC[11]. *GPNMB+* microglia show high expression of genes associated with lipid processing, cell migration, and vascular reorganization (Figure S3). Additionally, two other subgroups of microglia show clinicopathological associations consistent across all three population groups in the two cortical regions. In individuals with a clinical diagnosis of MCI, these include an over-representation of *CD74*-high microglia (with enrichment of mitochondrial process-associated genes) in STG and an under-representation of *CCL4+* cytokine-expressing microglia in DLPFC (Fig. 1F, S3).

### *SERPINH1+* and *CD44+* astrocytic subgroups show conserved disease associations across populations

After microglia, astrocyte clusters showed the largest number of conserved signatures associated with AD phenotypes, particularly in the caudate (Fig 1F). *SERPINH1+* astrocyte proportions were higher in the AC in individuals with higher neuritic plaque counts; this cluster is enriched in genes associated with lipid processing (Fig. S3). In addition, the AC-enriched astrocyte cluster expressing *WIF1* and *COX8A* was higher in individuals with a clinical diagnosis of MCI, suggesting a potential caudate-specific altered signature associated with. Additionally, *CD44+/KCNQ3+* astrocytes were less abundant in the AC in individuals with a clinical diagnosis of AD dementia; whereas cortical astrocytes expressing *CD44* have been classified as fibrous (as opposed to protoplasmic)[31], *CD44+* astrocytes in the caudate are less well studied, and their gene expression profile shows enrichment for processes associated with immune response signaling (Fig. S3). The lower abundance of this astrocyte cluster in individuals with AD dementia may point to a potentially protective role for these astrocytes in this brain region. In the cortex, *SERPINH1+* astrocytes were also more abundant in the STG in individuals with a clinical diagnosis of Alzheimer’s dementia. In the DLPFC, *WIF1+/VAV3+* astrocytes were more abundant in individuals at Braak stages 4-6; this astrocyte signature has been reported previously in DLPFC[11, 32], and may mediate tau-driven synaptic loss and cognitive decline. Overall, a wide range of astrocytic signatures were associated with AD phenotypes in a consistent manner across all three population groups in this study.

### Continuous signatures capture additional glial associations not found in discrete cluster analysis

Previous work by us and others[16, 33, 34] have suggested that the assignment of nuclei into discrete clusters may not capture continuous signatures that may have relevance to disease associations. This is particularly the case among glial cells, where distinct and potentially dynamic signatures within the same major class may exist without being specified through different developmental lineages. Using the scHPF workflow[35], we identified factors capturing continuous modes of correlated gene expression in each of our major cell classes. As an illustrative case, we identified 10 factors in astrocytes (Fig. 2A); some of these factors recapitulate the separation of nuclei into discrete clusters (e.g. Factor 8 is restricted to the *SERPINH1+* cluster), while others capture coordinated gene expression spanning multiple clusters (Fig. 2B,C, S4). Some of these factors highlight additional signatures that have associations with AD phenotypes but are not well captured by our assignment of nuclei into discrete clusters. This was particularly the case for oligodendrocytes: oligodendrocyte factor 4 (with high expression of *SLC5A3*, and enriched in genes associated with immune signaling and lipid processing, Fig. S5) is more prevalent in individuals with higher neuritic plaque counts, higher PHFtau tangle density, and a clinical diagnosis of Alzheimer’s dementia, in both the STG and the AC. By contrast, oligodendrocyte factors 3 (high expression of *LAMA2*, and enriched for genes associated with synaptic vesicle recycling) and 5 (high expression of *RABGEF1* and enriched for cytoskeleton-associated genes) were lower in both cortical regions in individuals with a clinical diagnosis of Alzheimer’s dementia (Fig. 2D,E, S5). These two oligodendrocyte factors are accompanied by lower expression of Astrocyte factor 6 (with expression of *SLC1A2*) in cortical tissue in individuals with a clinical diagnosis of Alzheimer’s dementia (Fig. 2D,E), thus providing additional candidate gene expression programs that are potentially depleted in cognitively impaired individuals. The discovery of these factors, which cross cluster boundaries in a non-trivial way, is thus complementary to the standard clustering approach described above. This lends further credence to examining both discrete and continuous models of gene expression variation when running disease associations, particularly in glia, where transcriptional programs may not be fully captured by discrete taxonomies.

**Figure 2:**
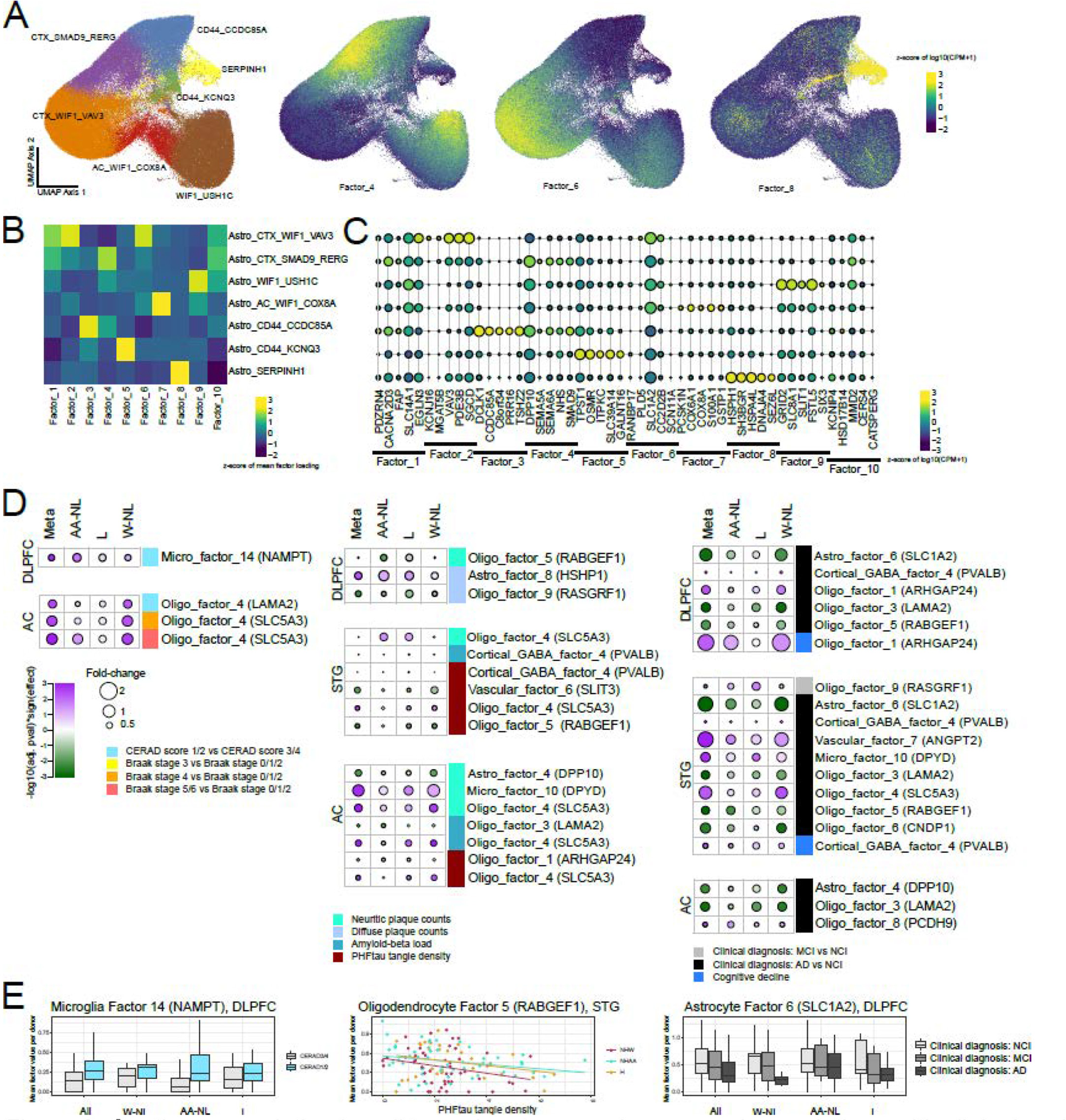
Continuous variation in cell type signatures and conserved associations with clinical and pathological traits across population groups. A) Visualization of scHPF-derived factor scores versus cluster identities in astrocytes. B) Heatmap showing median factor score in each cluster. Whereas some factor scores (e.g. Factor 8) are high in nuclei belonging to a specific cluster, others (e.g. Factors 1, 4 and 6) are represented in multiple clusters. C) Expression of genes with the top 5 weights in each factor, plotted by discrete cluster. Consistent with the representations in A and B, some factors are cluster-specific whereas others span multiple clusters. D) Effect sizes and p-values (from ANCOM-BC) of factor associations with consistent and significant (meta-analysis FDR-adjusted p-value <0.05 & I^2^<50%) signals in all population groups, in the same format as Fig. 1F. Unlike with the discrete clustering showing in Fig. 1, oligodendrocytes are highly represented in the factor-based analysis. E) Examples of factors with AD trait associations, in the same format as Fig. 1G.

### ATAC-seq analysis identifies putative regulators of cluster-specific signatures

Having identified key cell types and gene expression programs with significant and consistent associations across our population groups, we examined the ATAC-seq data to uncover putative drivers of cell signatures. Overall, our ATAC-seq data alone was not sufficient to identify robust subclusters within major cell types, nor did it yield robust statistically significant cell type-specific ATAC-seq peaks directly associated with clinical phenotypes across all three population groups; this is consistent with prior studies that have used the same technology, and sparseness of the data may be a result of relatively shallow sequencing depth (∼30k reads per nucleus) unable to overcome the inherent noise of the technique. By contrast, we did identify tens to hundreds of peaks that differed significantly across our transcriptomically-defined subclusters (Fig. 3A,B, S6), consistent with previous studies[22]. Sequence motif enrichment analysis of these differential peaks yielded putative transcription factor (TF) binding sites enriched in each cluster. Cross-referencing these with TF expression in the relevant cell types then yielded a smaller set of TFs that may drive some aspect of cluster identity (Fig. 3C). This putative driver set included *SREBF1* and *HSF1* in the *SERPINH1+* astrocyte cluster, *EGR1* in the *GPNMB+* microglia cluster, *WT1* in Astrocyte factor 6, *CREM* in Oligodendrocyte factor 3, *SIX3* in Oligodendrocyte factor 4, and *ELF4* in Oligodendrocyte factor 5.

**Figure 3:**
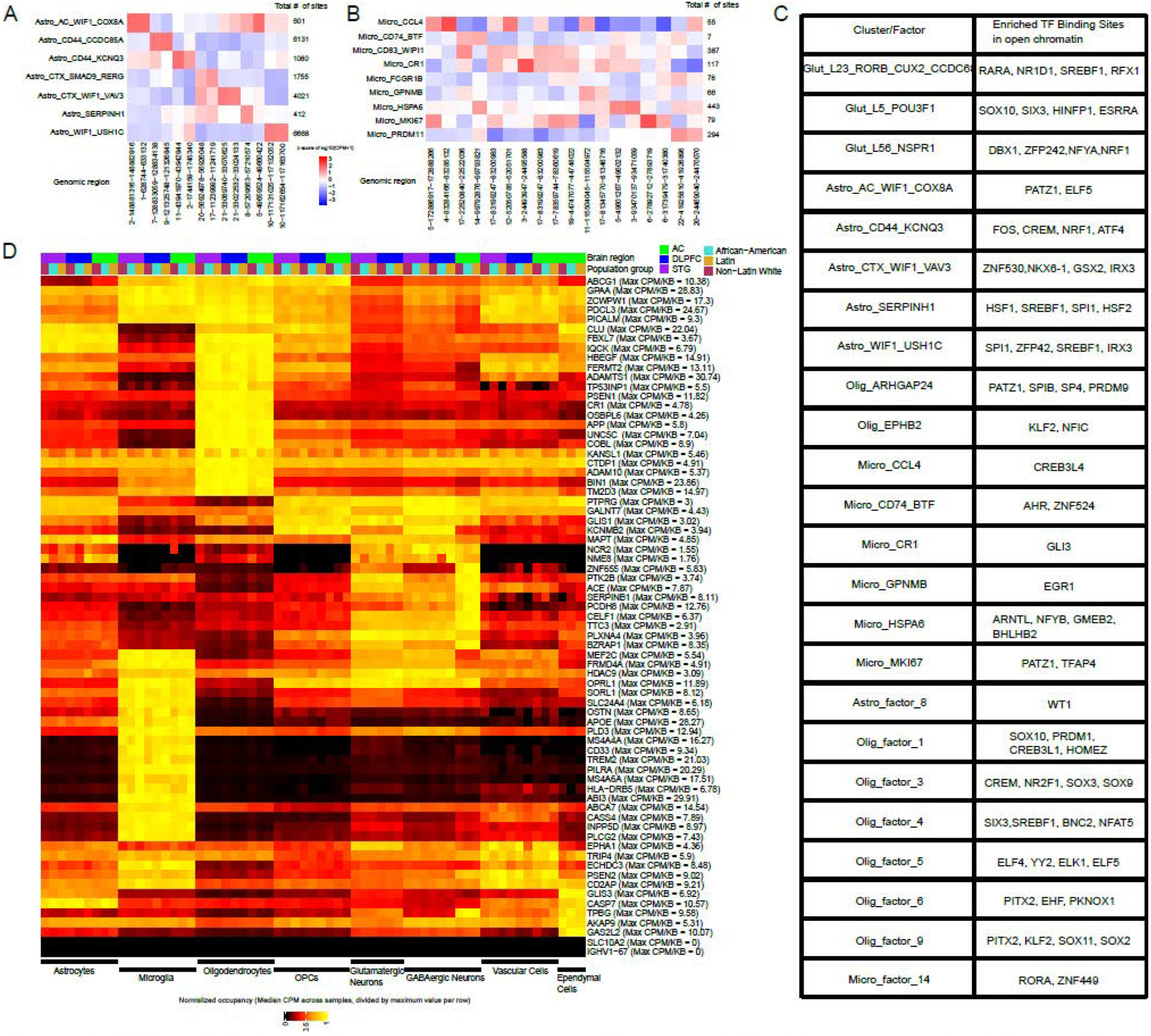
Differential chromatin occupancy across cell types and AD risk genes, and putative cell type drivers. A) Occupancy values (normalized counts per pseudobulked cluster) of top 2 differential sites (columns) for each astrocyte cluster (rows). Numbers on the right of the heatmap indicate the total number of differential sites per astrocyte cluster, in a one-versus-all comparison. B) Same as A, but for microglial clusters. C) Table of putative transcription factors whose binding sites show enrichment in the open chromatin regions for different clusters and factors. D) Normalized chromatin occupancy values at each of ∼70 AD-associated GWAS loci. Colors represent the total ATAC-seq counts over all nuclei for a given cell type, donor population group, and brain region, and then normalized to the maximum value for each locus across all donor/cell type/region combinations. Several loci show differences across cell types in occupancy, whereas only GABAergic neurons show strong region-specific differences, as expected based on their distinct regional expression signatures. Only one locus (containing the KANSL1 gene) shows consistent differences across the three participant population groups.

### Open chromatin regions near AD-associated genes are conserved in a cell type-specific way across population groups

Besides identifying putative drivers of cell signatures associated with AD, we also examined chromatin accessibility in each population at genetic loci implicated in AD through GWAS studies[36–51] (Fig. 3D). As expected, microglia show the highest accessibility at previously-described immune gene loci (including loci containing *TREM2*, *CD33*, and *HLA-DRB5*), consistent with cell type-specific eQTL studies highlighting these loci[5, 52–54]. Oligodendrocytes show the highest accessibility at many loci, including those containing early-onset AD genes such as *APP* and *PSEN1*, whereas neurons have the highest accessibility in loci containing *PLXNA4* and *NCR2*.

Despite the greater genetic diversity afforded by profiling individuals from multiple backgrounds, we only found one AD-associated locus with potentially differential occupancy across our population groups. This locus is on chromosome 17 (46,029,916 – 46,223,808 in GRCh38), proximal to the locus containing *MAPT*, and contains the *KANSL1* gene. It was identified in GWAS on predominantly White individuals[36] and on African-Americans[55, 56]. In our data, this locus is less accessible overall in African American individuals as compared to Non-Latin Whites (Fig. 3D). This difference in accessibility is consistent across all major cell classes but does not correspond to gene expression differences in *KANSL1* itself, suggesting that the effect of regulatory elements in this locus may act elsewhere in a population-specific manner.

### Compositional and expression signatures identify three molecular subgroups each of individuals with MCI and AD

Recent work has identified molecular subgroups of individuals based on bulk RNA-seq, bulk proteomics, and multi-modal examination. Here, we leveraged our multi-ethnic donor population to identify groups of molecularly similar tissue profiles across a wider range of genetic backgrounds and life experiences. Using a standard hierarchical clustering approach based on cluster and factor composition in STG tissue from individuals with MCI and AD, we identified six subgroups that maximized the silhouette score stability (Fig. 4A, Methods). Three of these subgroups were enriched for individuals with clinical diagnoses of MCI, whereas the other three each had >70% of individuals with clinical diagnoses of AD dementia. Whereas CERAD scores and Braak stages were not different across each of the three MCI-predominant subgroups and across each of the AD dementia-predominant subgroups, several clusters and factors distinguished the molecular subgroups. Among the three dementia-predominant subgroups, Group 4 was characterized by higher proportions in a subset of glutamatergic neuron clusters, as well as *ARHGAP24+* oligodendrocytes. Donors in group 5 had higher proportions of multiple glial signatures, including *CD44+* astrocytes and oligodendrocyte factor 9 (enriched for lipoprotein metabolism and receptor binding). Finally, group 6 was distinguished by higher expression of microglial factors 3,6,7, and 9 (enriched for genes involved in fatty acid transport, iron transport, phagocytosis, and lipoprotein processing, respectively), suggesting a molecular disease subtype associated with a subset of microglial signatures (Fig. 4A). Among the three AD dementia groups, Groups 4 and 5 are not particularly enriched for any race or ethnicity, whereas Group 6 (n=10) has an absolute majority of African American individuals (n=7), with a trend towards statistically significant enrichment (hypergeometric test with FDR-adjusted p-value = 0.081). Overall, this grouping suggests a complex landscape of molecular heterogeneity in individuals with dementia, which would merit further exploration with larger, representative sampling of the aged population. This work complements prior studies[3, 6, 11, 57] that have identified molecular subtypes using bulk and single-nucleus transcriptomics and proteomics on tissue, providing further insight into the molecular heterogeneity in tissue from individuals with Alzheimer’s pathology and dementia.

**Figure 4:**
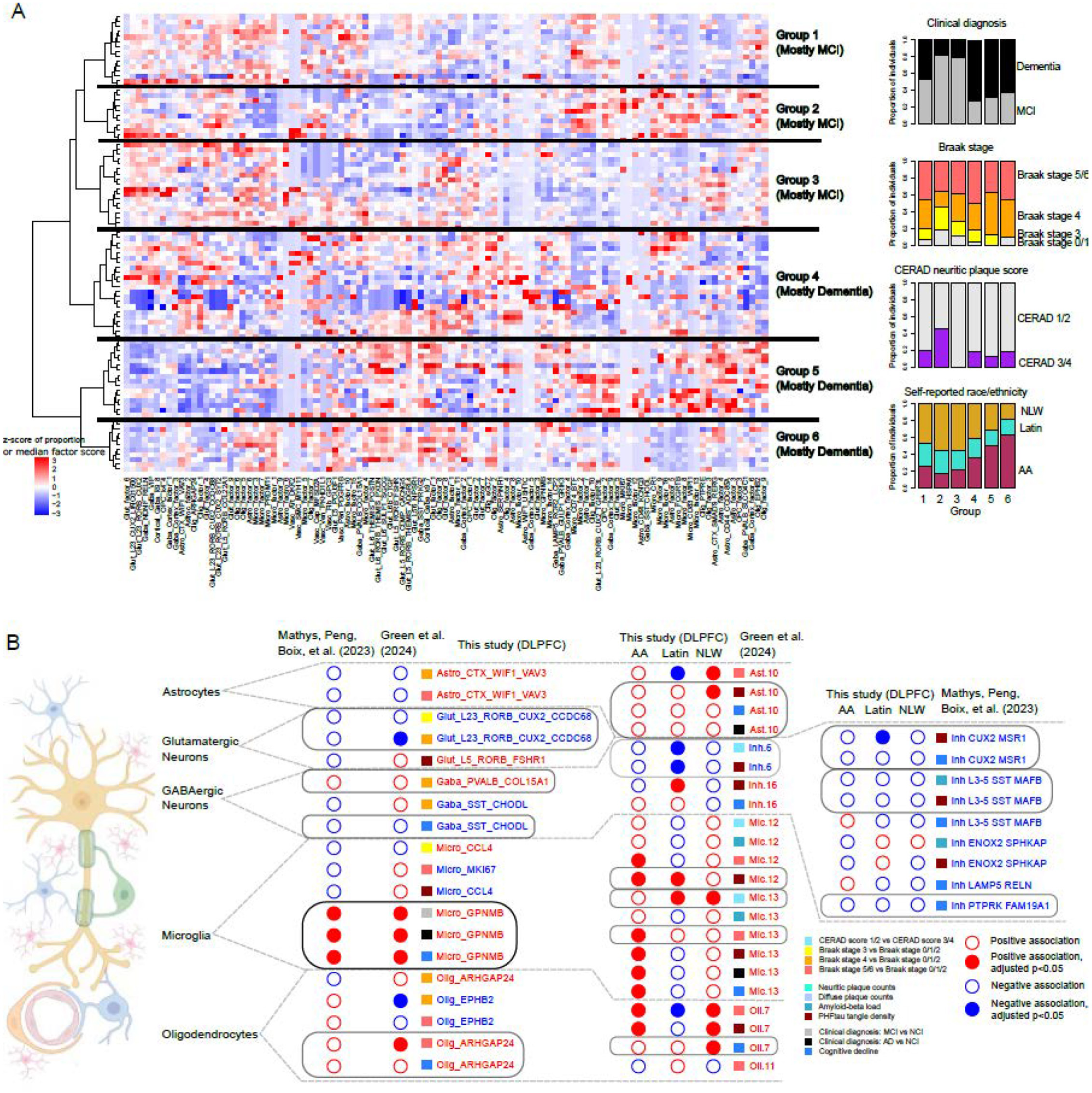
Molecularly-defined donor subgroups and replication of cell type changes in large-scale snRNA-seq studies. A) Heatmap showing 6 clusters of individuals (rows) based on relative proportions of cellular clusters and median factor values (columns). Bar plots on the right show the relative composition of each group based on terminal cognitive diagnoses, pathological staging, and population groups. B) Summary of cluster proportion findings that are replicated from this study in two other large-scale studies of predominantly White individuals from ROSMAP: Mathys, Peng, Boix et al. (2023) and Green et al. (2024). Colors indicate positive (red) or negative (blue) associations with clinical and pathological variables (see legend); solid circles represent FDR-adjusted p-values <0.05, whereas open circles indicate associations with adjusted p-values >0.05. Black boxes identify strongly replicated cell cluster findings (same direction of association and statistical significance), whereas grey boxes identify findings with similar directions of association and partial statistical significance. Left column shows results from mapping cluster labels from this study onto the other two studies. Middle column shows results from mapping cluster labels from Green et al. (2024) onto this study, with associations calculated separately across each population group. Right column shows results from mapping cluster labels from Mathys, Peng, Boix et al. (2023) onto this study. For the two external studies, only cluster labels reported as being statistically significantly associated with AD phenotypes.

### Agreement with prior studies focused on mostly White donors

To evaluate the robustness of our findings, we mapped our clusters to two large-scale single-nucleus RNA-seq studies on the DLPFC, both of which included primarily White individuals from ROSMAP[11, 20]. We found that *GPNMB+* microglia showed the same robust and statistically significant association (higher abundance) in the DLPFC in these two published studies, in individuals with a clinical diagnosis of MCI or Alzheimer’s dementia, and with steeper slope of cognitive decline (Fig. 4B). Other clusters, such as *ARHGAP24+* oligodendrocytes, were consistently higher in individuals with steeper slope of cognitive decline in both studies, though not reaching statistical significance (p<0.05). Similarly, *RORB+/CUX2+/CCDC68+* neurons were consistently lower in individuals at Braak stages 3 and 4, though statistically significantly so only in one study. The reverse mapping of cluster labels from Green et al.[11] also showed that Mic.12 and Mic.13, both of which express GPNMB and are putatively lipid-processing microglial subgroups, were more abundant in individuals at Braak stage 5 or 6, individuals with a clinical diagnosis of Alzheimer’s dementia, and individuals with higher PHFtau tangle density, in all three of our population groups; the same association was found with Oli.7, the analog of *ARHGAP24+* oligodendrocytes, and Ast.10, the analog of W*IF1+/VAV3+* astrocytes (Fig. 4B). Similarly, mapping of vulnerable GABAergic neuron clusters from Mathys, Peng, Boix et al.[20] to our data set showed that *SST+/MAFB+* neurons show a trend towards lower proportions in individuals with higher amyloid-beta load and PHFtau tangle density in all three of our population groups, and *CUX2/MSR1+* neurons show the same trend in individuals with higher PHFtau tangle density and steeper slope of cognitive decline (Fig. 4B). Thus, despite the relatively limited sample size per population group in our study, we find that some microglial, neuronal, and oligodendrocyte findings are consistent with findings from larger, though less diverse, studies.

## Discussion

In studies of aging and neurodegeneration, the inclusion of participants from various racial and ethnic backgrounds is important to answer two questions: 1) which disease-associated molecular signatures are conserved across different population groups, and 2) how do molecular subtypes of disease self-organize when including individuals with greater genetic heterogeneity and different life experiences? Here, generating single-nucleus RNA and ATAC-seq data from individuals of various backgrounds, we show that specific signatures in multiple cell classes – *GPNMB+* microglia, *SERPINH1+* astrocytes, a subset of *CUX2+/RORB+* glutamatergic neurons, and a range of oligodendrocytic factors – show similar associations with AD pathology and cognitive phentoypes in African-American, Non-Latin White, Latin individuals. In addition, with our broader sampling of the population, we also identify three molecular subgroups of individuals with dementia, each of which is distinguished by a subset of glial and neuronal signatures that complement the more universal conserved signatures described above. Thus, this study allowed us to ask and answer questions that were not addressable with prior single-nucleus data sets, and the results presented here prioritize specific cell clusters and signatures for pre-clinical studies that are applicable to a broader segment of the population.

A second key aspect of this study is the combination of discrete and continuous categorization of cellular heterogeneity within each major cell class. As has been reported by us and others before, continuous representation of certain glial classes (such as oligodendrocytes) may provide clearer associations with disease phenotypes, as compared to discrete cluster-based representation. By using both representations, we not only find a wider range of disease phenotype associations at the gene expression level, but also additional putative driver genes from the ATAC-seq data whose binding sites are enriched in clusters as well as in factors.

Despite these findings, there are two main caveats in this study, both having to do with statistical power. One of these is the relatively shallow depth of ATAC-sequencing per nucleus (median of 30k per nucleus); because the full space of accessible chromatin is much larger than the total length of DNA lying within gene boundaries, the variability and dropout of ATAC-seq signatures per nucleus is much higher. Indeed, our ATAC-seq data did not identify groupings of nuclei beyond what was found in the RNA-seq, and we did not find any conserved associations of individual chromatin regions that were associated with disease within a cell type across all three population groups after multiple testing correction. Future studies with substantially higher ATAC-seq depth may thus identify new disease-associated chromatin regions that are orthogonal to gene expression signatures.

The other main caveat in this study is sample size – although the total number of tissue samples processed (399) is on par with recent large-scale single-nucleus studies of AD[11, 20, 24], the total number of donors in each population group is less than 100. Thus, whereas this study is powered to ask about conserved signatures across all the population groups, it is less well-powered to ask about group-specific signatures, especially given confounders such as age of death (Fig. S1). Thus, findings like putatively differential chromatin accessibility at the AD-associated locus containing *KANSL1* (Fig. 3c), or the lack of cross-population group reproducibility of vulnerable neuronal signatures identified from prior studies (e.g. *LAMP5+/RELN+* GABAergic neurons, Fig. 4b, Supplemental Table 1) does not necessarily indicate race- or ethnicity-specific differences, but could rather be due to lack of power in the overall study design. Still, this lends weight to the conserved signature associations reported here, given that they are significant despite study confounds and power.

Finally, it is worth reiterating our use of self-reported race and ethnicity, as opposed to genetic ancestry, to define our population groups. Given that most of these donors lived in the United States from the 1940s and beyond, it is likely that their perceived (by themselves or by others) race and ethnicity impacted their life experiences. Thus, our analysis takes into account both genetic and life experience impacts on brain tissue signatures. By focusing on common signatures across the population groups, we prioritized specific cell signatures whose further exploration will likely have the broadest clinical impact. In parallel, our data also enables future studies that focus on transcriptomic and epigenetic associations with genetic variants; such studies aim to examine the specific effects of differing genetics on brain cell signatures in aged individuals. Thus, our work sets the stage for larger studies that consider this representative sampling approach of human genetic and life experience heterogeneity to understand the full phenomenology of aging and neurodegenerative diseases such as Alzheimer’s.

## Methods

### Study participants, ethics, and clinicopathological characterization

All brain tissue was obtained from participants in the Religious Order Study/Rush Memory and Aging Project (ROSMAP)[26], the Minority Aging Research Study (MARS)[27], and Latino Core[28] cohorts. As described previously, all participants are without known dementia at enrolment, have annual clinical evaluations; participants whose brain tissue was profiled in this study also consented to brain donation. At death, the brains undergo a quantitative neuropathologic assessment, and the participant’s rate of cognitive decline is calculated from the longitudinal cognitive measures that include up to 31 yearly evaluations[26]. An Institutional Review Board of Rush University Medical Center approved each study. All participants included in the analyses presented here signed an informed consent, Anatomic Gift Act and repository consent. For this study, we selected 164 participants, including all donors from the MARS, AA-Core (African American Core), and Latino Core studies who had full pathological characterization by December 2022, and availability of fresh frozen tissue from at least two of the three regions profiled (DLPFC, STG, or AC). As a result, our study cohort includes diverse individuals across the full range of the pathological stages and diagnosis of AD and mild cognitive impairment[58–60] A subset of demographic and clinicopathologic characteristics are summarized in Fig. 1 and Extended Data Fig. 1. Pathological measures were collected as previously described in Boyle et al.[61] and Yu, L. et al. [62]. We focused our analysis on the following measures:

1. CERAD neuritic plaque score[59, 63]: Semiquantitative measure of neuritic plaques. A neuropathologic diagnosis was made of no AD, possible AD, probable AD, or definite AD based on semiquantitative estimates of neuritic plaque density as recommended by the Consortium to Establish a Registry for Alzheimer’s Disease (CERAD), modified to be implemented without adjustment for age and clinical diagnosis. A CERAD neuropathologic diagnosis of AD required moderate (probable AD) or frequent neuritic plaques (definite AD) in one or more neocortical regions. Diagnosis includes algorithm and neuropathologist’s opinion, blinded to age and all clinical data. The coded values correspond to 1 (Definite), 2 (Probable), 3 (Possible), and 4 (No AD).
2. Braak stage[59]: Semiquantitative measure of severity of neurofibrillary tangle (NFT) pathology. Bielschowsky silver stain was used to visualize NFTs in the frontal, temporal, parietal, entorhinal cortex, and the hippocampus. Braak stages were based upon the distribution and severity of NFT pathology: Braak stages 1 and 2 indicate NFTs confined mainly to the entorhinal region of the brain Braak stages 3 and 4 indicate involvement of limbic regions such as the hippocampus Braak stages 5 and 6 indicate moderate to severe neocortical involvement. Diagnosis includes algorithm and neuropathologist’s opinion.
3. Neuritic plaque counts: A quantified measuredetermined by microscopic examination of silver-stained slides from 5 regions: midfrontal cortex, midtemporal cortex, inferior parietal cortex, entorhinal cortex, and hippocampus. The count of each region is scaled by dividing by the corresponding standard deviation. The 5 scaled regional measures are then averaged to obtain a summary measure for neuritic plaque counts.
4. Diffuse plaque counts: A quantitative measure determined by microscopic examination of silver-stained slides from 5 regions: midfrontal cortex, midtemporal cortex, inferior parietal cortex, entorhinal cortex, and hippocampus. The count of each region is scaled by dividing by the corresponding standard deviation. The 5 scaled regional measures are then averaged to obtain a summary measure for diffuse plaque counts.
5. Amyloid-beta load[64]: Quantified measure of Amyloid-beta protein identified by molecularly-specific immunohistochemistry and quantified by image analysis. The value is percent area of cortex occupied by amyloid-beta, and the overall score is the mean score in 8 regions (4 or more regions per individual are needed to calculate). The square root of this final score was used for association analyses
6. PHFtau tangle density[64]: Quantitative measure of neuronal neurofibrillary tangles, identified by molecularly specific immunohistochemistry (antibodies to abnormally phosphorylated Tau protein, AT8). Cortical density (per mm2) is determined using systematic sampling. Mean of tangle score in 8 regions (4 or more regions are needed to calculate). The square root of this final score was used for association analyses.
7. Clinical diagnosis[65]: Physician’s overall cognitive diagnostic category, based on all available clinical data at the time of death. All available clinical data were reviewed by a neurologist with expertise in dementia, and a summary diagnostic opinion was rendered regarding the most likely clinical diagnosis at the time of death. Summary diagnoses were made blinded to all postmortem data. Case conferences including one or more neurologists were used for consensus on selected cases. For this study, the categories used where NCI (No cognitive impairment – no impaired domain), MCI (Mild cognitive impairment – one impaired domain), and Alzheimer’s dementia.
8. Slope of cognitive decline: A quantitative measure based on uniform structured clinical evaluations, including a comprehensive cognitive assessment, are administered annually to the participants, extensively summarized in previous publications[66, 67]. Scores from 19 cognitive performance tests common in both studies, 17 of which were used to obtain a summary measure for global cognition as well as measures for five cognitive domains of episodic memory, visuospatial ability, perceptual speed, semantic memory and working memory. The summary measure for global cognition is calculated by averaging the standardized scores of the 17 tests, and the summary measure for each domain is calculated similarly by averaging the standardized scores of the tests specific to that domain. To obtain a measurement of cognitive decline, the annual global cognitive scores are modelled longitudinally with a mixed effects model, adjusting for age, sex and education, providing person-specific random slopes of decline (which we refer to as the slope of cognitive decline). For association analyses in this study, the negative of the slope value was used (so that higher values of the association variable correspond to steeper slopes of decline).

### Single-nucleus RNA+ATAC-seq (multiome) data generation

*Tissue preparation:* This study profiles post-mortem frozen human brain tissue isolated from Dorsolateral Prefrontal Cortex (BA9), Superior Temporal Gyrus (BA22), and Anterior Caudate, from individuals in the ROSMAP (Religious Orders Study, Memory and Aging Project) cohort, the MARS (Minority Aging Research Study) cohort, and Latino Core cohorts at Rush University. Tissue from each of these regions was dissected while frozen from flash frozen tissue blocks at Rush University and sent to Columbia University. Working on ice throughout, we carefully dissected to remove white matter and meninges, when present. The following steps were also conducted on ice: about 50-100mg of gray matter tissue was transferred into the dounce homogenizer (Sigma Cat No: D8938) with 2mL of NP40 Lysis Buffer [0.1% NP40, 10mM Tris, 146mM NaCl, 1mM CaCl2, 21mM MgCl2, 40U/mL of RNAse inhibitor (Takara: 2313B)]. Tissue was gently dounced while on ice 25 times with Pestle A followed by 25 times with Pestle B, then transferred to a 15mL conical tube. 3mL of PBS + 0.01% BSA (NEB B9000S) and 40U/mL of RNAse inhibitor were added for a final volume of 5mL and then immediately centrifuged with a swing bucket rotor at 500g for 5 mins at 4°C. Samples were processed 2 at a time, the supernatant was removed, and the pellets were set on ice to rest while processing the remaining tissues to complete a batch of 3 samples for each run of the 10x Genomics Chromium platform. The nuclei pellets were then resuspended in 500ml of PBS + 0.01% BSA and 40U/mL of RNAse inhibitor. Nuclei were filtered through 20um pre-separation filters (Miltenyi: 130-101-812) and counted using the Nexcelom Cellometer Vision and AO/PI stain at 1:1 dilution with cellometer cell counting chamber (Nexcelom CHT4-SD100-002).

*Library preparation and sequencing*: Approximately 20,000 nuclei from 3 samples (comprising different donor/region combinations) were pooled into a single sample, and these nuclei were run on the 10x Genomics Chromium platform using the 10x multiome protocol (Chromium Next GEM Single Cell Multiome ATAC + Gene Expression Reagent Bundle, PN-1000283) which can be accessed here. Briefly, following transposition GEMs were generated by combining barcoded Gel Beads, transposed nuclei, a Master Mix that includes reverse transcription (RT) reagents, and Partitioning Oil on a Chromium Next GEM Chip J (10× Genomics; PN-2000264). Incubation of the GEMs in a thermal cycler for 45 minutes at 37°C and for 30 minutes at 25°C generates full-length cDNA from poly-adenylated mRNA for gene expression (GEX) library and a Spacer sequence that enables barcode attachment to transposed DNA fragments for ATAC library. This was followed by a quenching step that stopped the reaction. Next, GEMs were broken, and pooled fractions were recovered. Silane magnetic beads were used to purify the first-strand cDNA from the post GEM-RT reaction mixture. Barcoded transposed DNA and barcoded full-length cDNA from poly-adenylated mRNA were pre-amplified by PCR and the products were used as input for both ATAC library construction and cDNA amplification for gene-expression library construction. Libraries were pooled and sequenced together on a NovaSeq 6000 with S4 flow cell (Illumina, San Diego, CA) at the New York Genome Center, for a target coverage of around 645 million reads per sample for RNA-seq and 645 million reads per sample for ATAC-seq. The scATAC-seq libraries were sequences as follows: Read 1N, 50 cycles; i7 Index, 8 cycles; i5 Index, 24 Cycles; Read 2N, 49 cycles. The snRNA-seq libraries were sequenced as follows: Read 1, 28 cycles; i7 Index, 10 cycles; i5 Index, 10 Cycles; Read 2, 90 cycles.

*Primary processing of sequencing data*: Raw fastq files were aligned to the genome/transcriptome and quantified using the CellRanger arc v2 package (https://support.10xgenomics.com/single-cell-multiome-atac-gex/software/pipelines/latest/using/count) with default parameters. RNA-seq counts files were then processed with the CellBender package[68] (https://github.com/broadinstitute/CellBender) to remove background signal, also with default parameters. The post-processed CellBender h5 files were then used for downstream analysis and demultiplexing.

*Demultiplexing of sample barcodes and assignment to donors*: Because each Multiome library consisted of nuclei from three samples from different donors, the original donor of nuclei contained in each droplet was inferred by harnessing SNPs in snRNAseq reads using the freemuxlet workflow in the popscle package[69] (http://github.com/statgen/popscle). Importantly, the pooling of samples was designed such that no pair of individuals was represented in more than one library – this design allowed us to demultiplex and assign nuclei to individuals even without their reference genotype information (multiplexing plan included in the data deposited, see Data Availability section below). Freemuxlet was run on each 10x Genomics run, with the assumption of 3 donors (each contributing one sample) in the mixture of nuclei. Finally, the donor-specific vcfs generated by freemuxlet were compared across all batches, allowing us to assign each donor group a unique identity based on the combination of batches it appears in. We then assigned each singlet nucleus to the original donor.

### Clustering of nuclei

In order to assign nuclei to individual clusters, we focused on the RNA-seq data, and clustered nuclei using the Seurat v4 package[30] (https://satijalab.org/seurat/) from each 10x Genomics batch separately to identify major cell classes – glutamatergic neurons, GABAergic neurons, oligodendrocytes, oligodendrocyte precursor cells, microglia, astrocytes, vascular cells, ependymal cells, and non-microglial immune cells. Only nuclei with >500 UMIs (based on RNA-seq) and mitochondrial UMI percentage <5% were retained. We then aggregated nuclei from each of these broad classes from all batches, and subsequently subclustered each broad class separately – this included 4 rounds where clusters with mixed signatures were iteratively removed. After data cleanup, nuclei from each broad cell type were clustered over a range of PCs and resolution parameter values, and selected an optimal clustering based on the Calinski-Harabasz and Davies-Bouldin scores. To assign nomenclature to each cluster, we examined the fraction of expression of each gene and each cluster, and selected the most differential genes, some in combination, to uniquely identify each cluster. The final parameters for each broad cell type were as follows: Astrocytes (10 PCs, resolution = 0.3); Ependymal cells (10PCs, resolution = 0.05); GABAergic neurons (10 PCs, resolution = 0.4); Glutamatergic neurons (10PCs, resolution = 0.4); Microglia (20PCs, resolution = 0.2); Oligodendrocytes (10 PCs, resolution = 0.05); OPCs (10 PCs, resolution = 0.05); Vascular cells (10PCs, resolution = 0.1). After each broad cell type was clustered, subclusters were merged if they did not have at least 4 genes with abs(log2-fold change)>2 in each direction for every pair of subclusters. For non-microglial immune cells, cells were not clustered, but rather mapped to the HuBMAP Azimuth annotation using the existing interface (https://azimuth.hubmapconsortium.org/). After QC, iterative clean-up, and removal of nuclei with ambiguous or doublet donor assignments, we retained 1,914,581 nuclei for downstream analysis, out of an initial set of 2,967,275 nuclei.

### Cell type proportion associations with clinical and pathological traits

To identify statistically significant differences in cluster proportions across different pathological and clinical conditions, we used two methods: one published method (ANCOM-BC[70]), and one quasibinomial analysis approach to assess cell cluster proportion differences, described in more detail below. P-values and effect sizes plotted in Figures 1 and 2 are all from ANCOM-BC, with the corresponding values for the quasibinomial model reported in Supplemental Table 1, For the quasibinomial model, we took advantage of the fact that relative cell type proportions are naturally distributed in the closed interval [0–1]. Therefore, we used logistic regression with a logit link to model response variables in that interval. However, compositional data tends to be enriched with zeroes, depleted of ones and left-skewed. We modeled this with a quasibinomial distribution that uses an extra parameter to account for overdispersion. We also accounted for heteroskedastic residuals using robust standard errors calculated with White’s original method using the R package sandwich. Using the R package performance, we assessed the model assumptions and the influence of outliers. We computed the 95% confidence intervals based on these robust standard errors using the R package broom. Finally, for each model, we ran multiple iterations randomizing the relative cell type proportions across participants to assess model calibration and the rate of false positives. For both methods, we only modeled cell types with at least one more observation than the total number of terms in the models being run: “proportion ∼ variable_of_interest + age_at_death + sex + pmi” for pathological traits and “proportion ∼ variable_of_interest + age_at_death + sex + pmi + education” for cognitive traits. Cell compositional analyses were done separately for each broad class: glutamatergic neurons (DLPFC and STG only), GABAergic neurons (including only the relevant GABAergic types in each region), microglia, astrocytes, oligodendrocytes, and vascular cells. OPCs were not tested, since they did not have any discrete subclusters, and non-microglial immune cells were also excluded, because of low cell numbers. Unless otherwise specified, we used the Benjamini-Hochberg method to correct for multiple hypotheses (across broad cell types, regions, and clinical and pathological traits) at a rate of 5%.

Finally, for the cell cluster association analysis in the DLPFC replication data sets[11, 20], we used ANCOM-BC on both the original reference cluster calls as well as the mapped cluster names, with the same associated variables and covariates as in our data set.

### Meta-analysis across population groups

To identify consistent signals across the three population groups, we modeled each group independently and then meta-analyzed. For all of our proportion analyses (ANCOM-BC and the quasibinomial model), we used the package metafor to run a random-effects meta-analysis using a restricted maximum likelihood estimation with the Hartung-Knapp-Sidik-Jonkman method to test individual coefficients. For cell types that did not converge with such default parameters, we implemented restarts increasing the coefficient estimation iterations and adjusting the step increase followed by using maximum likelihood or a fixed-effects model with a t-test for the coefficients. We assessed model performance through leave-one-out, outlier detection and inspection of the residuals. We determined that a specific cell type-phenotype association was conserved across all three population groups only if 1) the FDR-adjusted p-value <0.05, 2) the sign of association was the same in all groups, and 3) the meta-analysis I^2^ < 50%.

### Identifying cell type factors through Single-Cell Hierarchical Poisson Factorization

To identify factors modeling continuous gene expression variation, we used the single-cell hierarchical Poisson factorization method[35] to generate factors in each broad cell class. For GABAergic neurons, we calculated factors separately for the Anterior Caudate and the cortex (both regions aggregated). For glutamatergic neurons, we only calculated factors for the cortex, given the paucity of glutamatergic neurons in the caudate. For each run of scHPF on each broad cell class, we identified the optimal number of factors by selecting the largest factor set such that no two factors had a gene-weight vector or cell-weight vector correlation greater than 0.75. For selection of genes to represent each factor for visualization, we selected the five genes with the largest weights, as well as a mean pseudobulked counts per million >10 across all nuclei in the broad class of interest.

For factor associations with AD phenotypes, we calculated the median factor value per donor+region combination, and ran simple linear models with the same general covariates as in the proportion analyses described above: “proportion ∼ variable_of_interest + age_at_death + sex + pmi” for pathological phenotypes and “proportion ∼ variable_of_interest + age_at_death + sex + pmi + education” for phenotypes that include assessments of cognition.

### Gene Ontology analysis

For enrichment analysis, differential genes were identified for each subcluster versus all other subclusters of the same major cell class using the standard edgeR package workflow, with counts pseudobulked per individual sample (donor + region). Genes were first filtered to only include those with at least 10 counts per million in at least 10 pseudobulked samples for the cluster of interest. Genes for enrichment were selected based on the following criteria: 1) Benjamini-Hochberg False Discovery Rate-adjusted p-value <0.05, 2) Mean pseudobulk counts per million >=10 for the cluster of interest, and 3) at least 2-fold higher expression (log2FC>1) in the cluster of interest. Genes satisfying this criterion were then assessed for “Biological Process” enrichment using the topGO R package, filtered to those processes with a Fisher’s test adjusted p-value <0.1, and aggregated into broader categories using the rrvrgo package.

For factors, genes were filtered to include only those detected in 5% of pseudobulked samples in a given cell class, and with a mean counts per million across all pseudobulked samples >5. Of these genes, the top 100 were selected for each factor based on their loading scores, and were fed directly into the topGO package for “Biological Process” enrichment. Aggregation and visualization was done in the same way as for cell clusters, using the rrvrgo package with pathways selected based on a Fisher’s test adjusted p-value <0.1.

### ATAC-seq visualization and differential peak analysis

ATAC-seq peaks were first calculated separately for each batch using Cellranger arc v2, as described above. Peaks were then aggregated separately across major RNA-defined cell classes (Glutamatergic neurons, GABAergic neurons, Astrocytes, Oligodendrocytes, OPCs, Vascular Cells, Microglia, and Other Immune Cells) after the RNA QC steps outlined above; *importantly, this resulted in a different set of peak regions called for each major cell class*. Peak aggregation and harmonization across samples within each cell class was done using the Signac library in Seurat. Clustering of nuclei within each major cell class using ATAC-seq data was performed within each RNA-defined major cell class using the standard workflow in Signac (Latent Semantic Indexing for dimensionality reduction, followed by Louvain community detection), but did not identify robust subclusters based on the CH and DB indices, so no ATAC-seq subclusters were called. UMAP visualizations for ATAC-seq data and for joint RNA- and ATAC-seq data were performed on a subset of 300,000 nuclei randomly sampled across all populations; peaks were harmonized across all nuclei regardless of major cells class. UMAPs for RNA-seq were based on PCA for dimensionality reduction (30 PCs used), UMAPs for ATAC-seq were based on LSI for dimensionality reduction after harmonizing peak calls across the 300,000 nuclei, and UMAPs for joint RNA- and ATAC-seq were based on joint clustering using the WNN workflow in the Signac[71] R package.

Differential ATAC-seq occupancy among RNA-defined subclusters was performed using the DESeq2[72] R package on pseudobulked ATAC-seq data. Pseudobulk counts were generated by summing counts for all peak regions within each donor+region sample for that subcluster, after harmonization of peaks within each major cell class. DESeq2 was run for each subcluster (against pseudobulked samples from all other subclusters within the same major cell class) with included batch, donor post-mortem interval, and donor age of death as covariates. All p-values were adjusted using the Benjamini-Hochberg False Discovery Rate correction. For visualizing the top 2 peaks per subcluster (Fig. 3A), peaks were sorted by p-value, and the top 2 peaks with positive log2-fold change were selected.

For Transcription Factor Binding Site and motif enrichment analysis (Fig. 3B), all cluster-specific differential peaks with FDR-adjusted p-value <0.05 and positive log2-fold changes were input into the XSTREME algorithm in the MEME suite using the interactive submission mode (https://meme-suite.org/meme/doc/xstreme.html). TFs with enriched binding sites in each subcluster were then selected after filtering out sequence motifs with >8 repetitions of the same nucleotide and filtering out sequences found in more than 1 subcluster.

### Open chromatin quantification for AD risk loci

ATAC-seq peak counts were pseudobulked as described above, but for all nuclei from a donor+region sample belonging to each major cell class (instead of each subcluster, as in the description above). Pseuodbulked values were then normalized to Counts Per Million within each donor+region+cell class sample. For each AD risk loci, CPM counts for all peaks overlapping with the locus (even partially, at the start and end points of the locus) were summed together for each sample. To allow for relative differences in peak locations and widths across different major cell class, these summed CPM values were then divided by the total length of peak regions, and then the mean length-normalized CPM value was calculated for each population group for all samples of a given region and major cell class. Finally, for plotting, the population mean length-normalized CPM value for each locus was divided by the maximum value (across all major cell classes and population groups). The values displayed in Fig 3c, therefore underwent four transformations: 1) sample-specific normalization for total read count, 2) length-normalization to account for total peak length within a locus, 3) mean calculation/aggregation for each of our three population groups, 4) normalization to the maximum value for a given locus, to allow for visualization of loci simultaneously.

### Mapping cell cluster labels across studies

Cluster calls from this data set were mapped to nuclei from the Green et al. and Mathys et al.[73] data sets using the mapmycells package from the Allen Institute for Brain Science (https://portal.brain-map.org/atlases-and-data/bkp/mapmycells). Nuclei from this study were randomly assigned to 5 groups, and a separate mapmycells model was built for each set of nuclei. The nuclei from the query data sets[11, 20] were then assigned 5 cluster labels, one from each of the 5 models. Nuclei that received the same label from at least 4 of the 5 models were assigned that cluster label, and nuclei that did not meet this criterion were called “ambiguous” and not retained for downstream analysis. This same procedure was also run in the other direction, for mapping cluster names from these two data sets (as reference) onto this study (as query).

### Donor clustering based on cell subcluster frequencies and median factor scores

To identify potential subgroups of donors based on cell type profiles, the within-broad class normalized proportions of astrocyte, neuronal, oligodendrocyte, microglial, and vascular clusters, as well as the median factor values per donor, were concatenated into input vectors for hierarchical clustering approach, with 1 – Pearson’s correlation r-value as the distance measure and Ward’s method for linkage. The resulting hierarchical tree was cut at various heights, and silhouette scores calculated at each height. The median silhouette score at each cut height was used to determine that 6 groupings was optimal. For each of these 6 groups, the number of donors for each cognitive diagnosis, Braak stage, CERAD score, and population group were calculated, as shown in Figure 4a.

## Data availability

All raw (fastq files), batch processing/multiplexing plan (flat file) and processed data (post-QC counts tables and cluster identities) are available on the AD Knowledge Portal, under the following DOI: https://doi.org/10.7303/syn53649093.

## Funding acknowledgements

This work was funded by the Foundation for the National Institutes of Health (to V.M.) through the Accelerating Medicines Partnership-Alzheimer’s Disease 2.0 (AMP® AD 2.0), as well as NIH grants R01AG066381 (to V.M.), R01AG22018 (to L.L.B.), P30AG072975 (to Julie Schneider), P30AG10161, R01AG15819, R01AG17917, U01AG46152, and U01AG61356 (to D.A.B.).

AMP® AD 2.0 is a public-private partnership managed by the Foundation for the National Institutes of Health and funded by the National Institute on Aging (NIA) in partnership with the private sector. For up-to-date information on the partners, visit https://fnih.org/our-programs/accelerating-medicines-partnership-amp/amp-alzheimers-disease-2-0/. ACCELERATING MEDICINES PARTNERSHIP and AMP are registered service marks of the U.S. Department of Health and Human Services.

## Supplemental Tables & Figures

**Supplemental Table 1**

Effect sizes and p-values of ANCOM-BC and quasibinomial models to assess proportion differences associated with clinical and pathological phenotypes. Columns represent individual population groups as well as the meta-analysis across all three groups (see Methods).

**Supplemental Figure 1.**
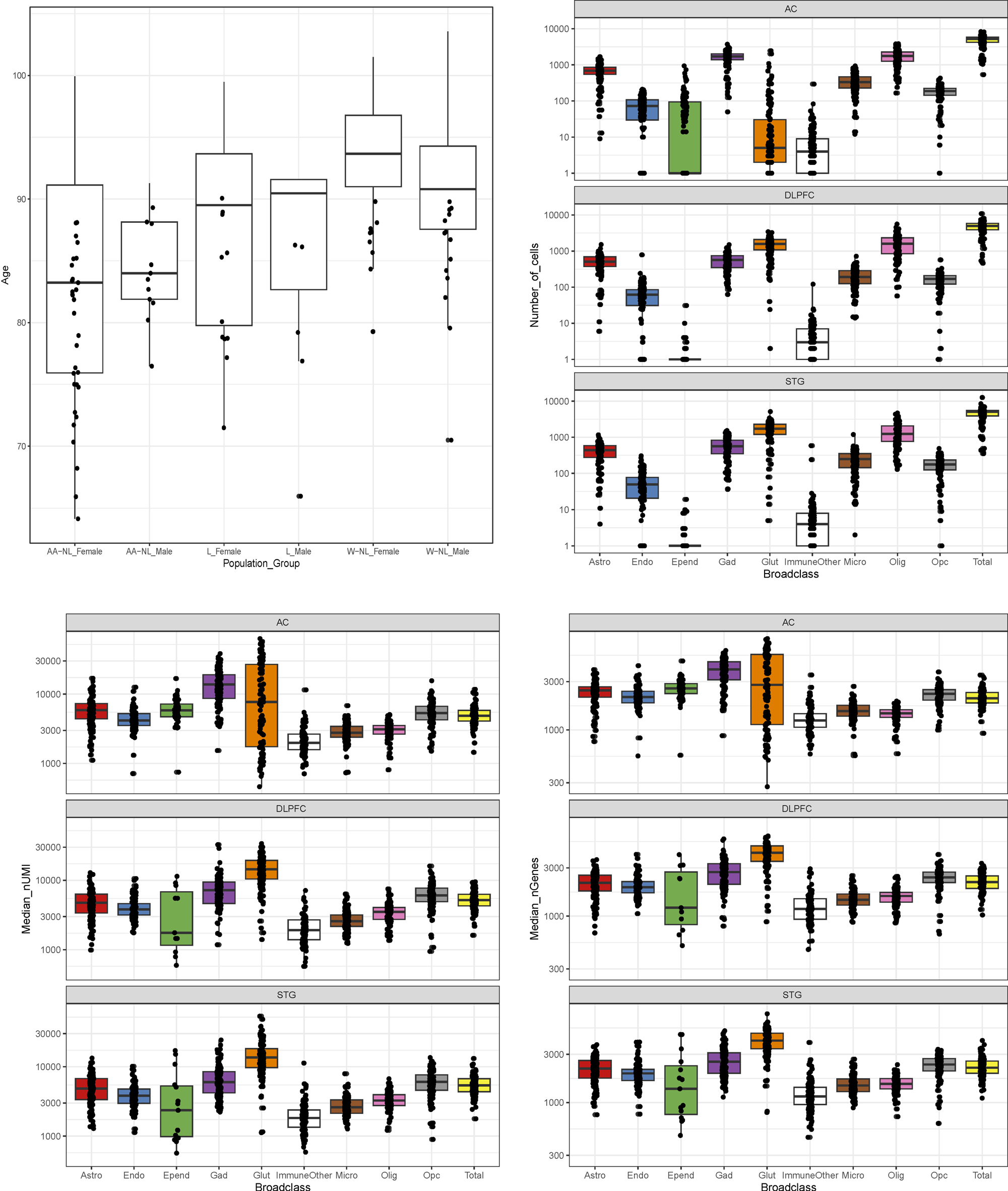
Distributions of age (top left) across donor groups, nuclei numbers per cell type per region (top right), total gene counts per nucleus per cell type per region (bottom left), and total transcript (Unique Molecular Identifier) counts per nucleus per cell type per region. For age, the boxplot distribution includes all participants whose brains were profiled, but individual points for those older than 90 have been removed from the visualization.

**Supplemental Figure 2.**
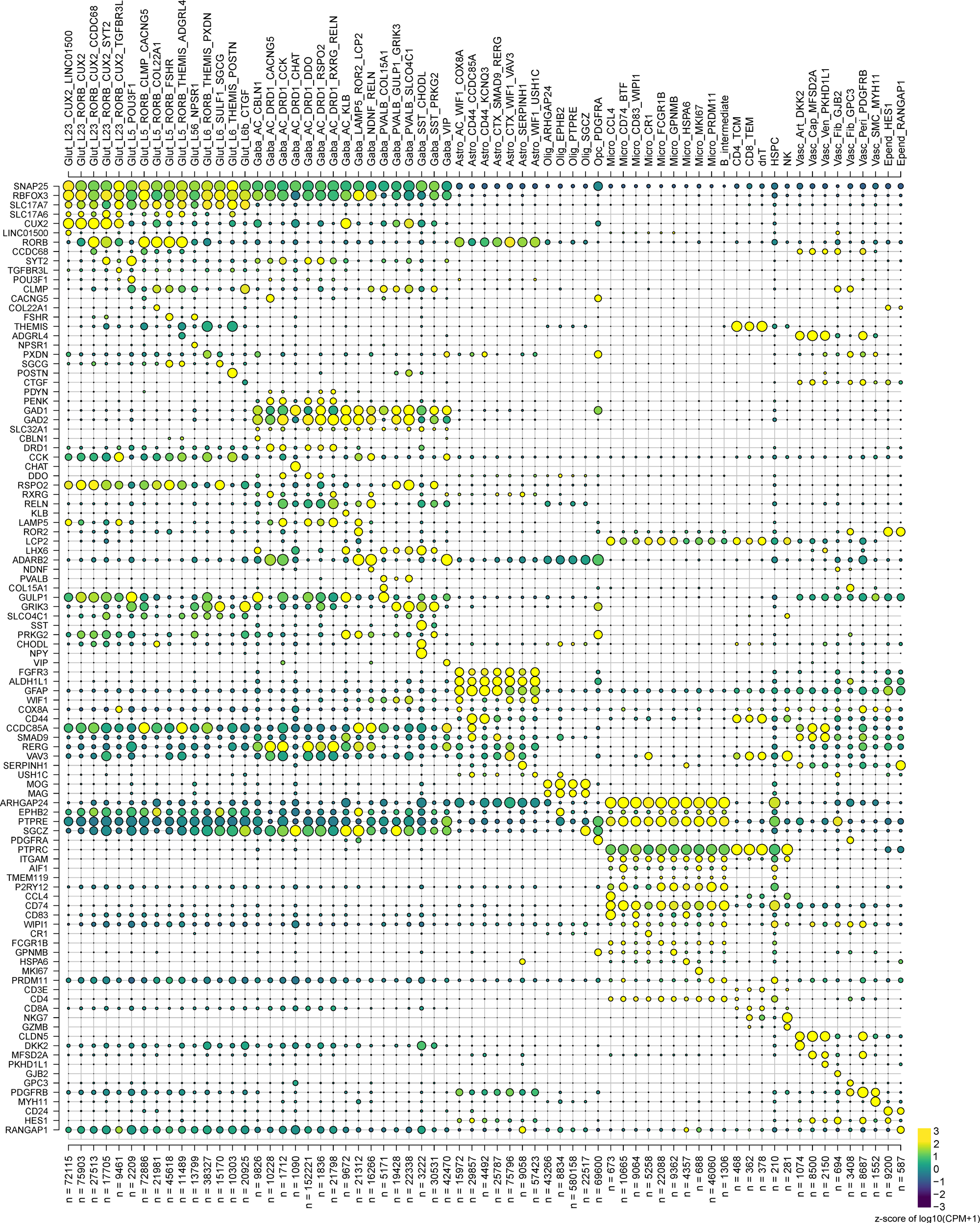
Dotplot showing the fraction of nuclei and mean gene expression in each cluster of broad cell class markers as well as key distinguishing genes between subclusters within each broad class.

**Supplemental Figure 3.**
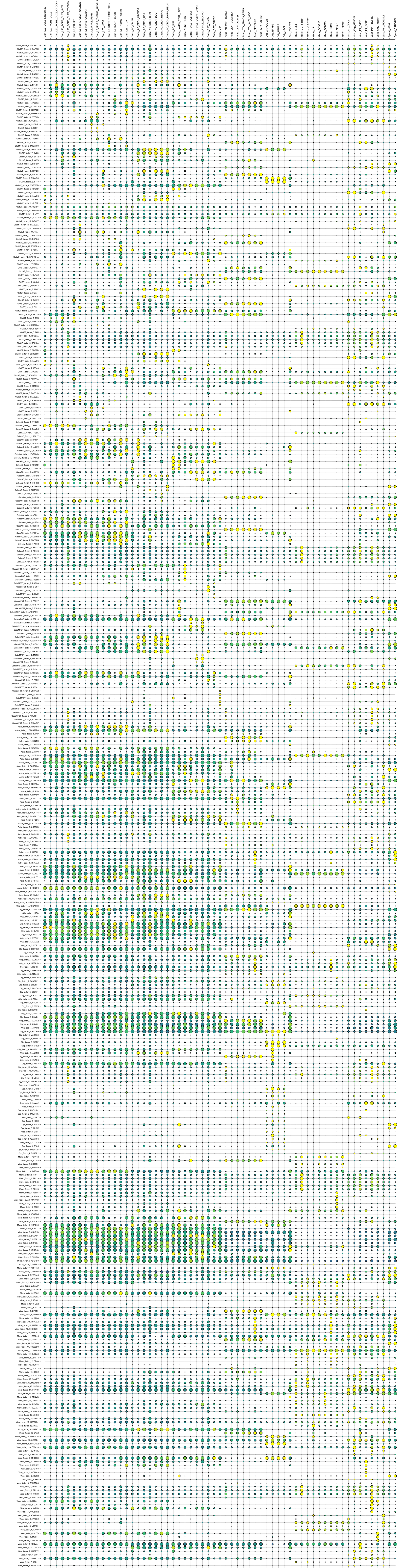
Compilation of Gene Ontology (GO) terms and aggregated categories for genes differentially expressed within each subcluster versus all other subclusters in the same main class (see Methods).

**Supplemental Figure 4.**
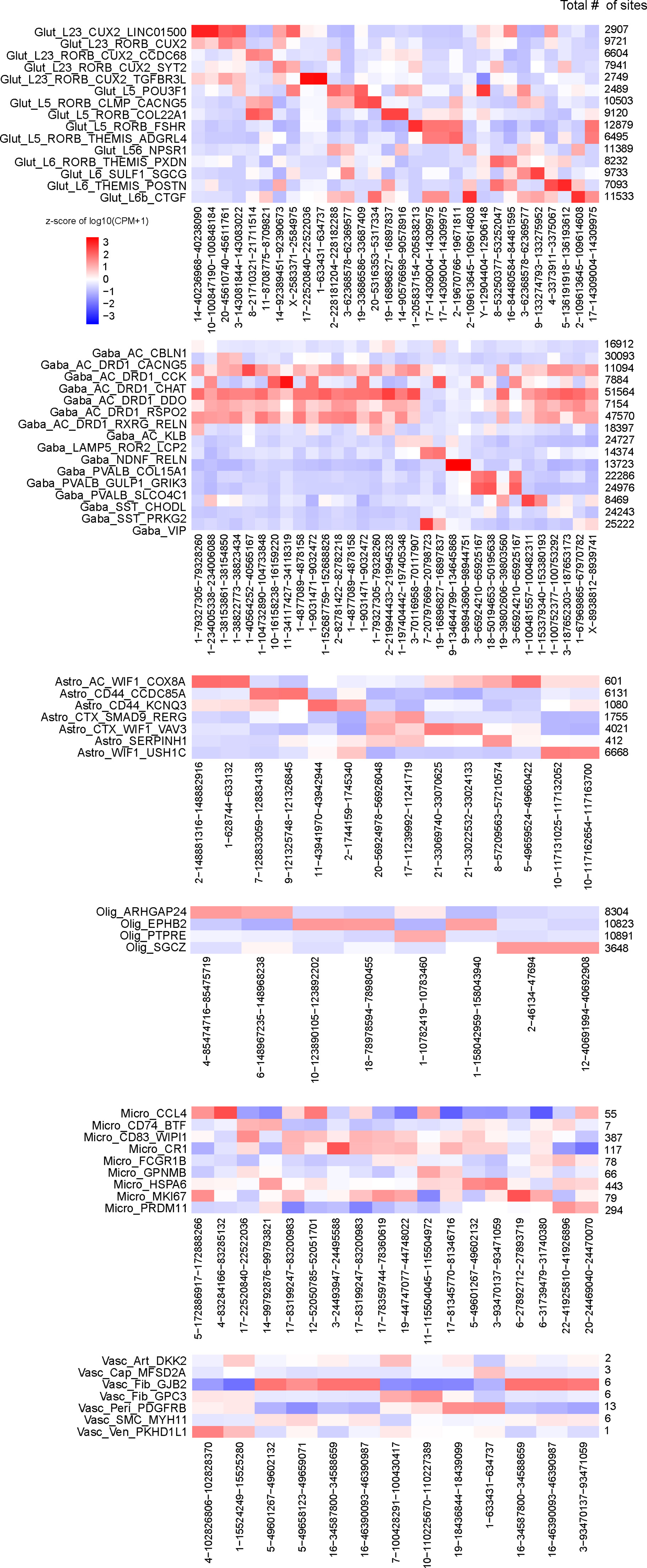
Dotplot showing the fraction of nuclei and mean gene expression in each cluster of the top 5 genes (by loading) on each scHPF-derived factor (see Methods).

**Supplemental Figure 5.**
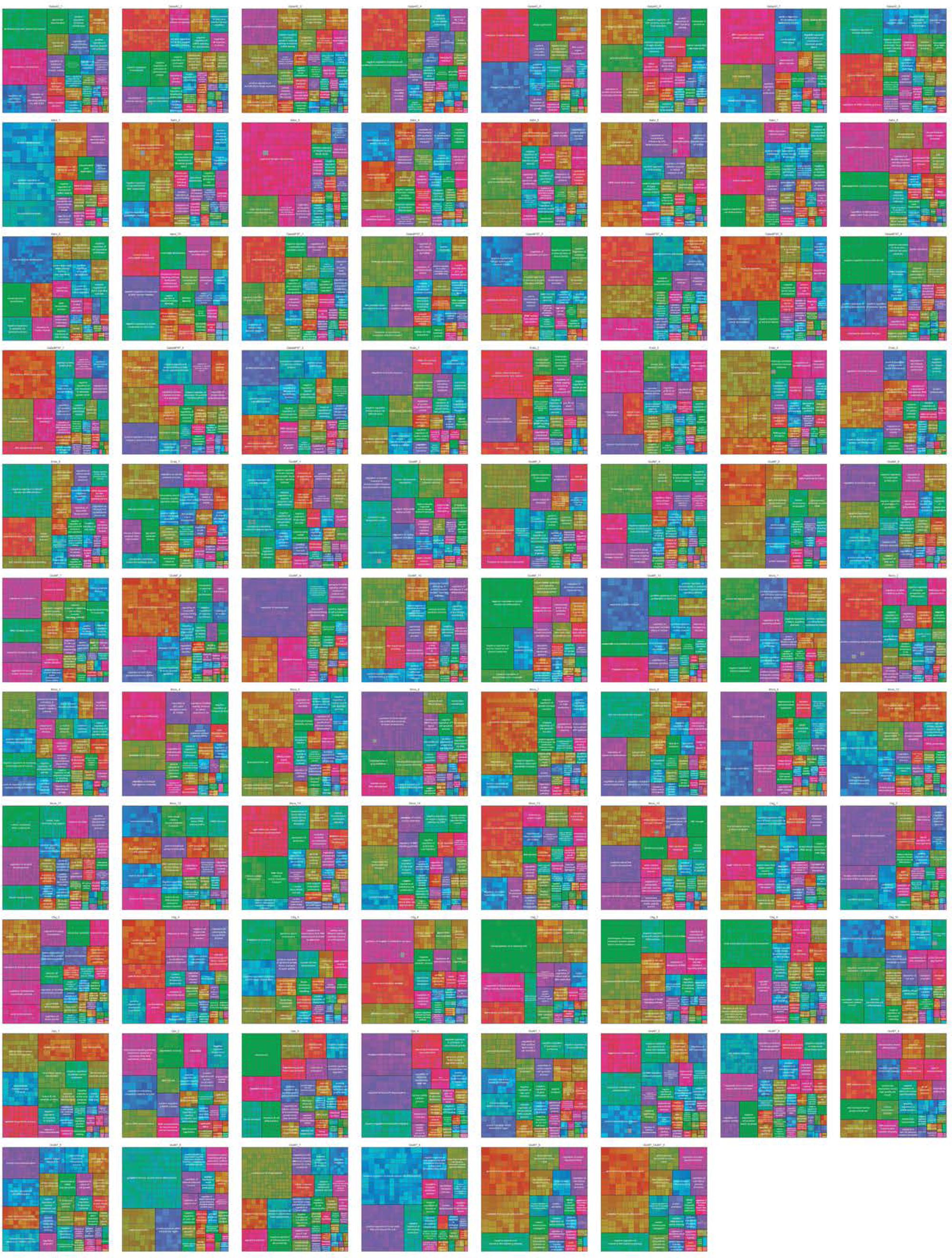
Compilation of Gene Ontology (GO) terms and aggregated categories for genes with highest loadings on each factor (see Methods).

**Supplemental Figure 6.**
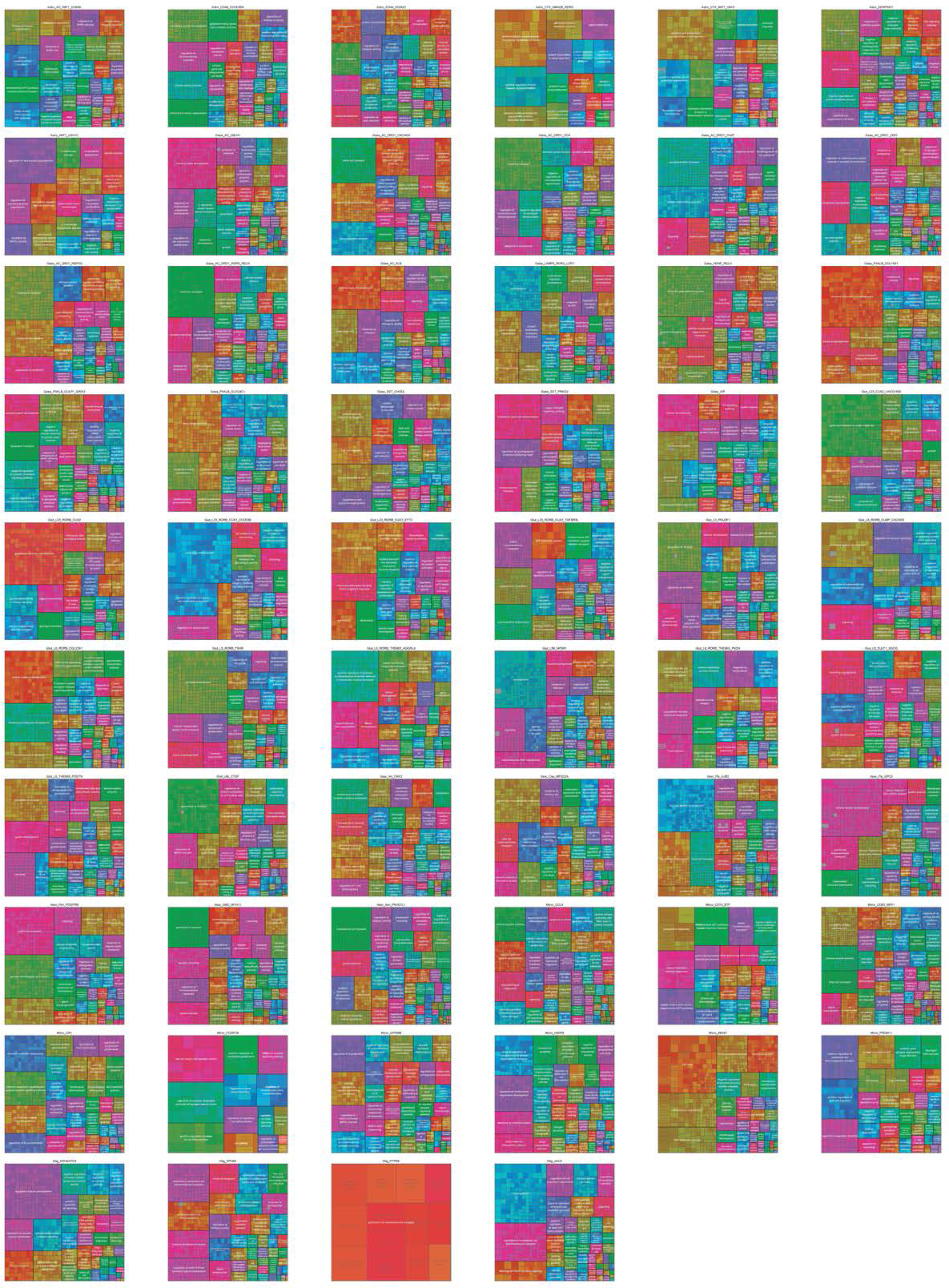
ATAC-seq-based occupancy for key cluster-enriched peaks, with clusters defined based on RNA-seq; this is analogous to Main Figure 3A and 3B. For each major cell class, the heatmap represents the z-score of the mean peak counts per million (pseudobulked over all nuclei of each cluster per sample). For each cluster, the top 2 differential peaks (versus all other clusters of that cell class) are included. Numbers on the right of each heatmap indicate the total # of differential sites per cluster.

